# Injury-­Induced Remodeling of Junctional Actin Bands in the Vestibular Maculae of Mice and Chicks: Implications for Sensory Regeneration

**DOI:** 10.1101/2025.11.24.682866

**Authors:** Mark E. Warchol

## Abstract

The vestibular organs of birds are capable of regenerating sensory hair cells after ototoxic injury, but the regenerative ability of the mammalian vestibular organs is much more limited. The factors that inhibit regeneration in the mammalian inner ear are not known, but it has been proposed that the structure of filamentous actin cables at cell-cell junctions within the sensory epithelium may be an important regulatory influence. Junctional actin cables in the chick utricle are relatively thin, while those in mouse utricle are much thicker. These differences result in differing mechanical properties of the avian vs. mammalian inner ear, which may affect the potential for regenerative proliferation. The present study characterized injury-evoked changes in junctional actin cables in the utricles of mice and chicks. We found that the thickness of junctional cables in the chick utricle was not affected by ototoxic injury, but that injury to the mouse utricle led to the formation of many new junctional actin bands whose thickness was comparable to those in the chick utricle. Thicker actin bands persisted after injury, but were not necessarily associated with cellular junctions. In addition, the relative extent of supporting cell expansion in the injured chick utricle was larger than that in the mouse utricle, which may affect activation of Hippo/YAP signaling in both species. Together, these data point to important differences in actin cable plasticity in the avian vs. mammalian utricle that may partially account for their differing regenerative abilities.

## Introduction

The sensory hair cells of the vestibular organs detect head movements and the orientation of the head with respect to gravity. Vestibular hair cells can be damaged or lost following ototoxicity or as a consequence of normal aging. Nonmammalian vertebrates can regenerate vestibular hair cells after ototoxic injury, leading to functional recovery. This process of hair cell regeneration has been particularly well-studied in the inner ears of birds and occurs largely (but not exclusively) by the renewed proliferation of supporting cells, which divide to produce new hair cells and supporting cells [1]. In contrast, the vestibular organs of mammals have a very limited ability to regenerate hair cells [2]. Mammalian vestibular organs can produce a moderate number of new hair cells after severe injury, but those replacement cells do not fully differentiate and do not restore sensory function [3,4]. Ototoxic damage to the mammalian vestibular organs also results in minimal cell proliferation [5,6]. Together, these observations suggest a link between the ability of supporting cells to divide in response to hair cell injury and the potential for functional hair cell regeneration.

Most studies of vestibular regeneration in birds and mammals have focused on the utricle, a sensory organ that detects linear acceleration. The factors that are permissive for regenerative proliferation in the avian utricle and that limit such proliferation mammalian utricle are not fully understood, but recent work has pointed to a key role for mechanical signals in regulating cell cycle entry. Cell-cell junctions in all vestibular epithelia possess circumferential actin cables, which link a cell’s cytoskeleton to the adherins junctions between adjoining cells. In the vestibular organs of birds, these junctional actin bands are relatively thin (∼0.5 µm), while those in mammals are considerably wider (∼3 µm) [7]. Junctional actin bands in the mammalian utricle increase in thickness during embryonic development, and this increase is correlated with reduced proliferative ability [7,8,9]. These observations suggest that the structure of junctional actin bands and the mechanical environment of the sensory epithelium may limit regenerative potential.

Mechanical regulation of cell division is mediated by the YAP signaling pathway, which can be activated by cellular stretching [10]. Prior studies have shown that YAP signaling regulates proliferation in the developing and regenerating utricle [11,12,13]. The lumenal surfaces of supporting cells undergo rapid expansion after hair cell loss, in order reseal the fluid barrier between endolymph and perilymph [14]. It is possible that this expansion process may also initiate YAP signaling, but the relative degree of supporting cell expansion in birds vs. mammals has not been quantified. To better understand the mechanical factors that may influence regeneration in the utricle, the present study quantified changes in the width of circumferential actin belts, as well as the spreading of supporting cells in the utricles of chicks and mice after ototoxic injury. We found that ototoxic injury did not affect the thickness of circumferential actin bands in the chick utricle, but that hair cell loss did lead to the formation of many thin actin bands in the mouse utricle. In addition, hair cell loss caused a greater relative degree of cellular expansion in the chick utricle than in the mouse utricle. These results have implications for the differential activation of YAP signaling in the vestibular organs of these species.

## Methods

### Animals

Studies used mice of both sexes and either129S1/SvImJ:C57Bl/6J background (JAX #101045). Chickens (White Leghorn strain) were hatched from fertile eggs (Charles River SPAFAS) and maintained in heated brooders. Chicks and mice had *ad libitum* access to food and water and were housed in the animal facilities of Washington University in Saint Louis. All protocols involving animals were approved by Institutional Animal Care and Use Committee (IACUC) of Washington University, School of Medicine, in Saint Louis, MO.

#### Ototoxic Lesions

Mice (129S1/SvImJ:C57Bl/6J, age 2-4 months) received a single i.p. injection of 3,3’-iminodipropanenitrile (IDPN, TCI America, mixed 1:1 in sterile PBS) at a dose of 4 mg/gm. Mice were maintained for an additional 3-56 days after IDPN injections. Chickens (2-4 weeks post-hatch) received injections of streptomycin sulfate (1200 mg/kg, i.m.) once/day for three consecutive days. Chickens were allowed to recover for 24 h after the last injection. All animals were euthanized by CO_2_ inhalation and isolated temporal bones (mice) or utricles (chicks) were immediately removed and fixed in for 1 hr 4% paraformaldehyde in 0.1 M phosphate buffer.

#### Immunohistochemistry

After fixation, utricles were thoroughly rinsed in 0.01M phosphate buffered saline (PBS, Sigma) and then processed for immunohistochemistry as wholemounts or as frozen sections. Specimens were first incubated for 2 hr in PBS with 5% normal horse serum and 0.2% Triton X-100 (Sigma). They were then incubated overnight (at room temperature) in primary antibody solution. The following primary antibodies were used: anti-Myosin-VIIa antibody (rabbit polyclonal, Proteus BioSciences, 1:500), and anti-β-catenin (mouse monoclonal, BD Biosciences, 1:100). All antibodies were diluted in PBS with 2% horse serum and 0.2% Triton X-100). The next day, specimens were rinsed 3X (>5 min each) in PBS and incubated for 2 hr in secondary antibodies (conjugated to Alexa-488, Alexa-568, and Alexa 647, Life Technologies, 1:500 in PBS with 0.2% Triton X-100) at room temperature. Filamentous actin was labeled with Alexa Fluor 555-conjugated Phalloidin (Invitrogen), and cell nuclei were labeled with DAPI (Sigma, 1 μg/ml). Finally, all samples were rinsed 3× (>5 min each) in PBS and mounted in glycerol:PBS (9:1) on glass slides.

#### Imaging and quantification

Specimens were imaged using a Zeiss LSM700 confocal microscope. Images were processed and analyzed using Volocity or ImageJ/FIJI Software. All quantification was conducted using high magnification images that were obtained using a 63x 1.4 NA objective. Hair cell counts were obtained from four 2,500 µm^2^ regions (two in the striola, two in the extrastriola). ImageJ/Fiji software was used to measure junctional thickness from high magnification images of phalloidin-stained specimens. The thickness of actin cables at junctions between adjoining supporting cells was measured, beginning at the left side of a high magnification image and moving rightward (see tracings in Fig 2A-C). The thickness of junctions between supporting cells and hair cells was not measured. Figures were assembled using Adobe Illustrator. *Statistical analysis*. Data analysis and statistics were carried out using GraphPad Prism version 6.0d. Data are presented as mean±SD. Student’s t-tests or analyses of variance (ANOVA) followed by Tukey’s or Bonferroni’s post hoc tests were applied, as appropriate.

## Results

### (I) Remodeling of junctional actin bands after ototoxic injury

Cellular junctions in the sensory epithelia of the inner ear possess filamentous actin bands that surround the inner perimeter of each cell and are linked to actin bands in adjoining cells via adherins junctions [8]. The thickness of these actin bands varies among the ears of different vertebrates and quantification of actin band thickness in vestibular maculae from five vertebrate classes suggests that the thick junctional actin cables in the mammalian ear may limit its regenerative abilities [8,19]. Motivated by the study of Burns et al. [7], we first used phalloidin to label filamentous actin in the utricles of mice and chicks. We found that the mean thickness of junctional bands in the mouse utricle was 2.88 ± 0.86 µm (n=279 samples from seven specimens, ages: P28-52), while the mean thickness of junctional bands in the chick utricle was 0.40 ± 0.08 µm, n=239 samples from five specimens, ages:10-14 days post-hatch). No differences in actin band thickness between the striolar and extrastriolar regions were noted in either species. These data indicate that junctional actin bands in the mouse utricle are ∼6-7x as thick as those in the chick utricle and are in agreement with the observations of Burns et al. [7].

We next characterized changes in the junctional actin bands in response to ototoxic injury. Most prior studies of ototoxicity have focused on hair cell damage caused by treatment with aminoglycoside antibiotics or cisplatin. However, cisplatin causes damage to actin structures in the inner ear [16] and the vestibular organs of mice are relatively insensitive to aminoglycosides [17]. Instead, we lesioned mouse vestibular hair cells via systemic treatment with 3,3’-iminodipropanenitrile (IDPN) [18,19]. Mice (2-4 months of age) received a single 4 mg/gm i.p. injection of IDPN and were allowed to recover for 3, 5, 7 or 56 days. At these time points, utricles were removed, fixed and processed for histochemical labeling of hair cells and actin filaments. Images of utricles from IDPN-treated mice revealed a reduction in hair cell density in the central region of the utricle, when compared to undamaged controls (Fig. 1, A-E). Hair cell density was quantified in 2,500 µm^2^ regions located within the striolar and extrastriolar regions of each utricle at 3, 5, 7 or 56 days after IDPN injection. These data revealed a progressive loss of hair cells in both regions over a seven day period. Consistent with other studies [19], we also observed modest hair cell recovery in both regions after 56 days recovery (Fig. 1F, G, p<0.0001).

**Figure 1.**
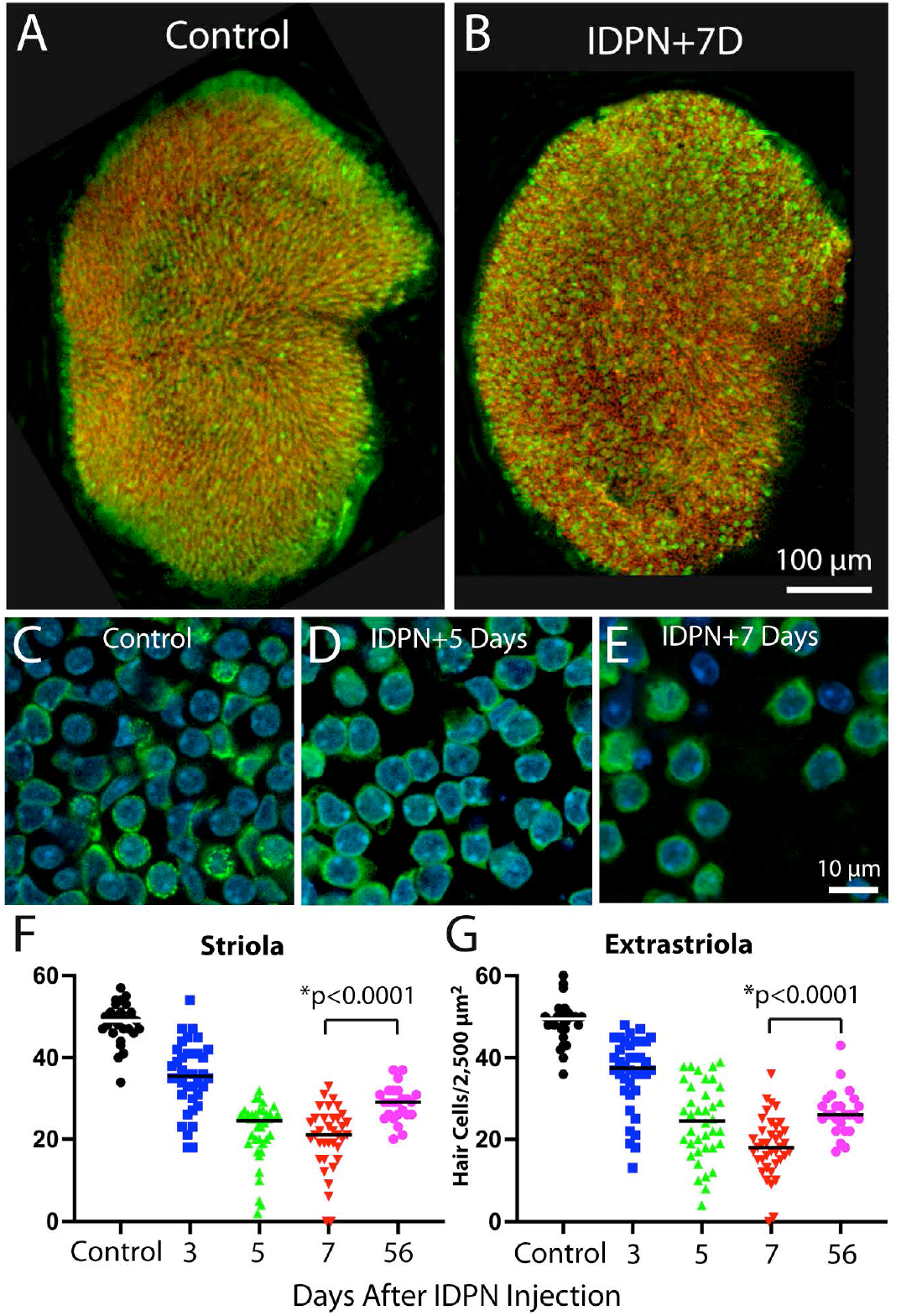
Ototoxic injury to the mouse utricle caused by systemic treatment with IDPN. Mice received a single 4 mg/gm injection of IDPN and were allowed to recover for 3-56 days. (A, B): At seven days post-lDPN, reduced hair cell density was observed in low magnification images of whole mounts. (C, D, E): Higher magnification images show hair cell loss. Such images were used to Quantify hair cell density at various recovery times. (F, G): IDPN injection led to a progressive decrease in hair cell density in both the striolar and extrastriolar regions of the utricle. Note that a small degree of hair cell recovery was observed in both regions after 56 days recovery. Labels: green=myosin VIia (hair cells), red=phalloidin (actin filaments), blue=DAPI (cell nuclei).

Having characterized IDPN-induced hair cell loss, we then examined resulting changes in the junctional actin belts. We found that ototoxic injury resulted in the formation of junctional actin bands with considerably reduced thickness, similar to the changes reported by Kaur et al. [20] following diphtheria toxin-induced hair cell loss in Pou4f3-huDTR transgenic mice. Utricles fixed at seven days after IDPN treatment had a mean junctional thickness of 1.20 ± 1.01 µm (n=274 measurements from five specimens; Fig. 2, B, C), Many of these junctions appeared to form ‘rosettes’ or ‘scars’ that were suggestive of epithelial repair after hair cell loss (Fig. 2E) [14]. Utricles fixed at 56 days post-IDPN had a mean junctional thickness of 3.22 ± 4.53 (n=211 samples from five specimens; Fig. 2C, D). These data indicate that the injured utricle possesses many thin actin belts, but that normal junctional thickness is restored after sufficient recovery time.

**Figure 2.**
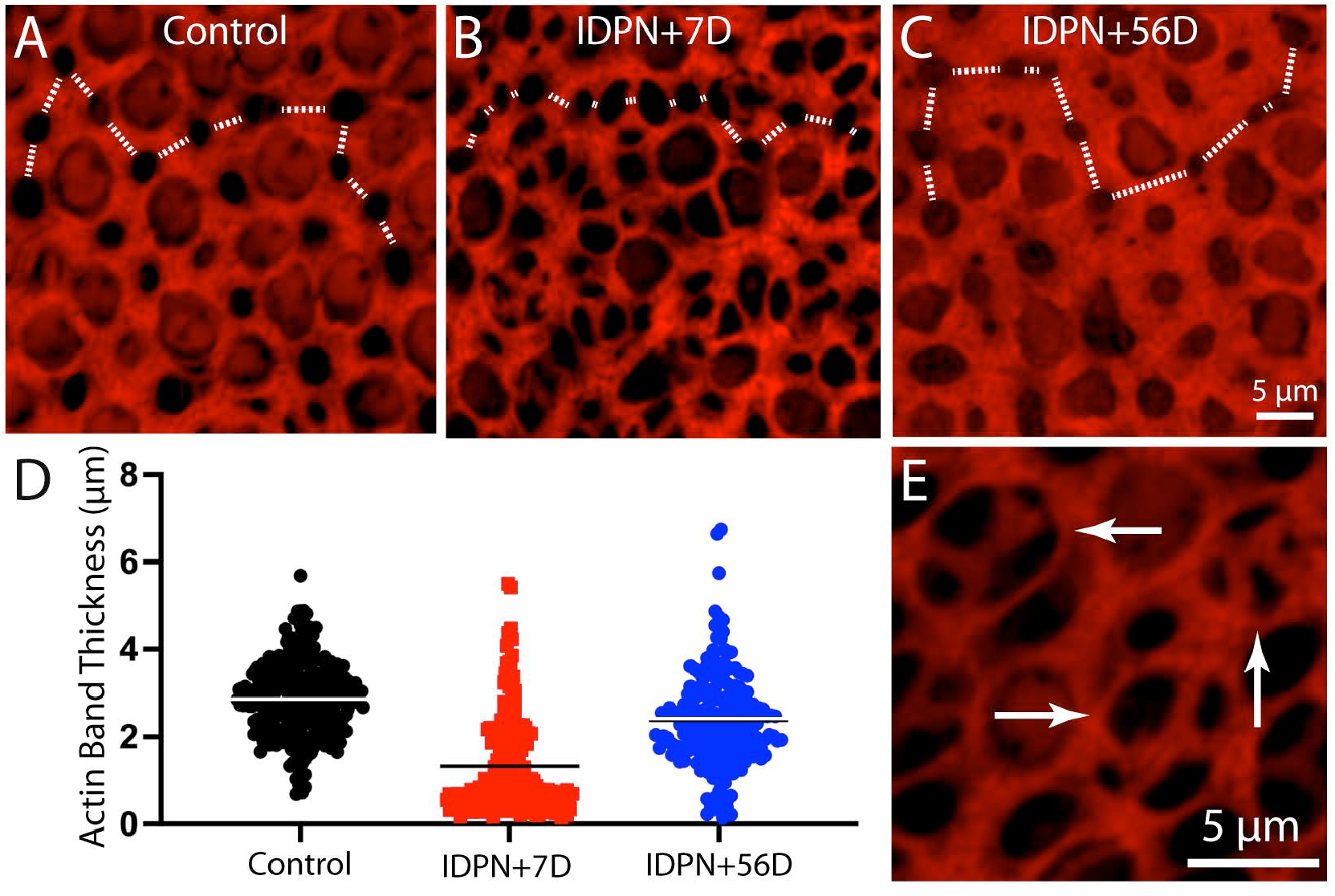
Changes in the thickness of junctional actin bands in the mouse utricle after IDPN-induced injury. The width of phalloidin-labeled junctional bands between supporting cells was quantified, moving from left to right across high magnification images (dotted white lines in A, B, C). (D): Quantitative data indicate that ototoxic injury results in a significant reduction in actin band thickness after seven days recovery. After 56 days recovery, the thickness of actin bands had nearly recovered to its original dimensions. (E): Many of the thin actin bands form ‘rosettes or ‘scars’ that mediate resealing of the sensory epithelium (arrows). Color: red=phalloidin (actin filaments).

Since the dimensions of junctional actin bands may influence the regenerative properties of supporting cells, it was of interest to compare actin bands in the utricles of mice vs. chicks after ototoxic injury. Hair cell lesions were created in chicks (2-4 weeks post-hatch, n=4) by giving three injections of 1,200 mg/kg streptomycin sulfate (one/day for three days) and then allowing then to recover for 24 hr after the final injection. [Note: Data on injury-induced nuclear translocation of YAP1 in some of these specimens has been published elsewhere - see ref. 13.] Fixed utricles were immunolabeled for myosin VIIa (to label hair cells) and phalloidin (to label actin junctions). Prior studies have shown that hair cell loss from this streptomycin regimen is limited to the striolar region of the utricle [13], so all data were obtained from the striola. Hair cell density in the striolar region of undamaged chick utricles was 53.6+6.9 HCs/2,500 µm^2^ (n=40 samples from five specimens), which decreased to 32.7+9.6 HCs/2,000 µm^2^ after streptomycin treatment (n=21 samples from eight specimens). The thickness of junctional actin bands in the utricles of streptomycin-treated chicks was 0.38+07 µm (n=202 measurements from eight specimens; Fig. 3B, C), which is nearly identical to the value reported above for uninjured utricles. Notably, plotting junctional thickness data obtained from injured mouse utricles on the same coordinates as data from chicks revealed that ototoxic injury caused many actin junctions in the mouse utricle to acquire similar thicknesses to those observed in the utricles of chicks (Fig. 3C).

**Figure 3.**
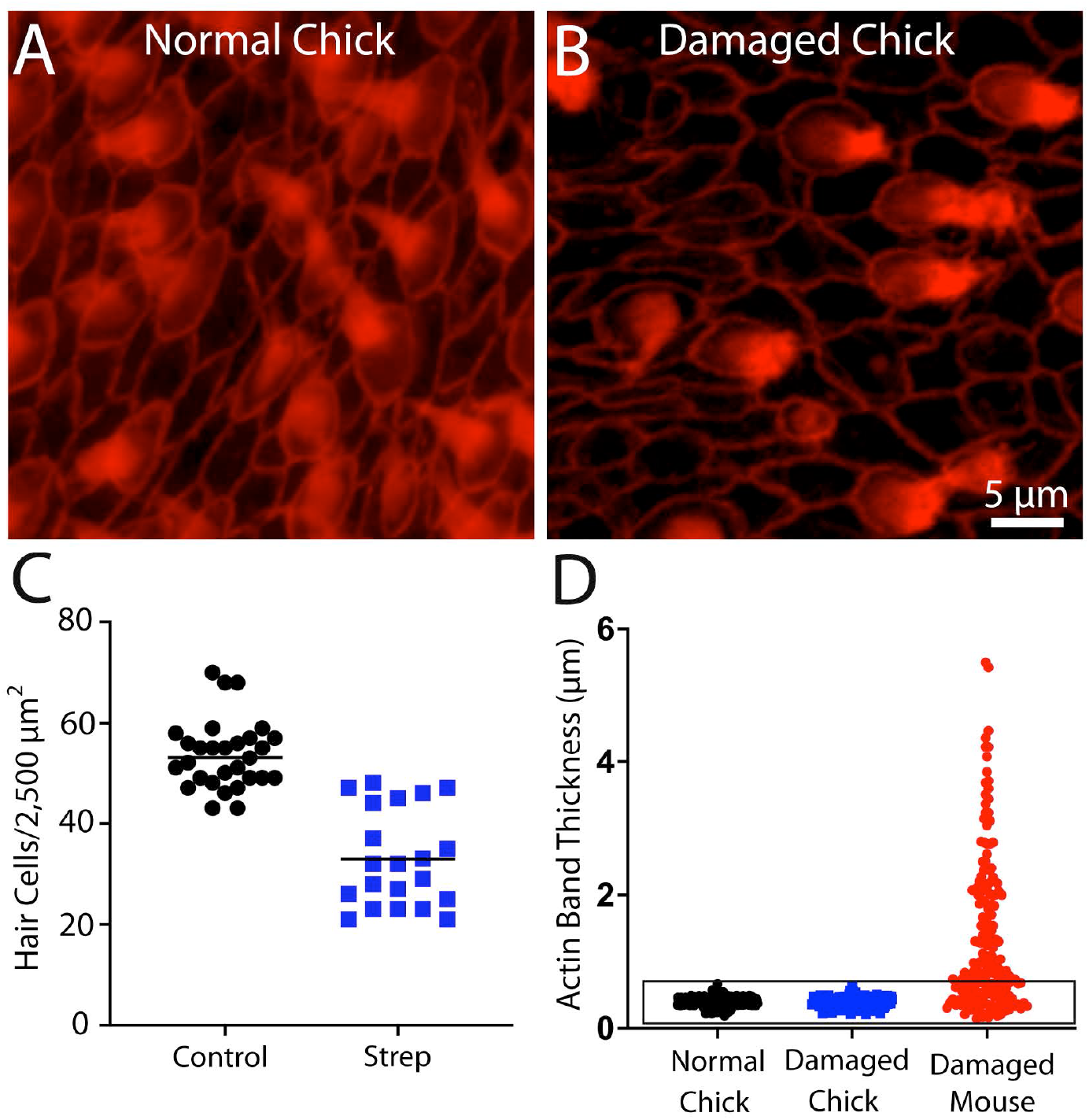
Ototoxic injury to the chick utricle does not affect the thickness of circumferential actin cables. Chicks received three daily infections of 1,200 mg/kg streptomycin sulfate and utricles were fixed 24 hr after the final injection. (A, B): Images of phalloidin-labeled (red) cell-cell junctions in the normal and streptomycin-damaged chick utricle. (C): Streptomycin treatment caused ∼60% reduction in hair cell density within the striolar region. (D): Quantification of actin band thickness from the striolar regions revealed no difference between normal and streptomycin-damaged chick utricles. Note, however, that potting data from the mouse utricle obtained at seven days after IDPN injection (shown in Fig. 2) on these same coordinates indicates that the thickness of many actin bands in the injured mouse utricle is comparable to those present in the chick utricle (D, data enclosed in box).

The loss of hair cells will create gaps in the lumen of the sensory epithelium, leading to disruption of the barrier between endolymph and perilymph. As noted earlier, such injury sites are quickly resealed by the formation of ‘scars’, which form when portions of 3-5 adjoining supporting cells extend into the epithelial space that was formally occupied by the lost hair cell [14]. In the present study, such structures were commonly observed in utricles fixed at 3-7 days after IDPN injection (e.g., Fig. 2E). They consisted of thin cables within the injured regions that were likely formed from actin processes that extended from supporting cells and joining near the center of the lesion. This combination of thick and thin actin cables at the lumen made it difficult to definitively identify cell-cell junctions in surface views of the injured sensory epithelium. Instead, adherins junction were identified by immunoreactivity for β-catenin [21]. As expected, β-catenin labeling in undamaged utricles overlapped with the circumferential actin bands (Fig. 4A, A’, A’’). However, in IDPN-treated utricles, a subset of the thicker actin bands did not co-label for β-catenin (Fig. 4B,C ), suggesting that they were no longer associated with cell-cell junctions.

**Figure 4.**
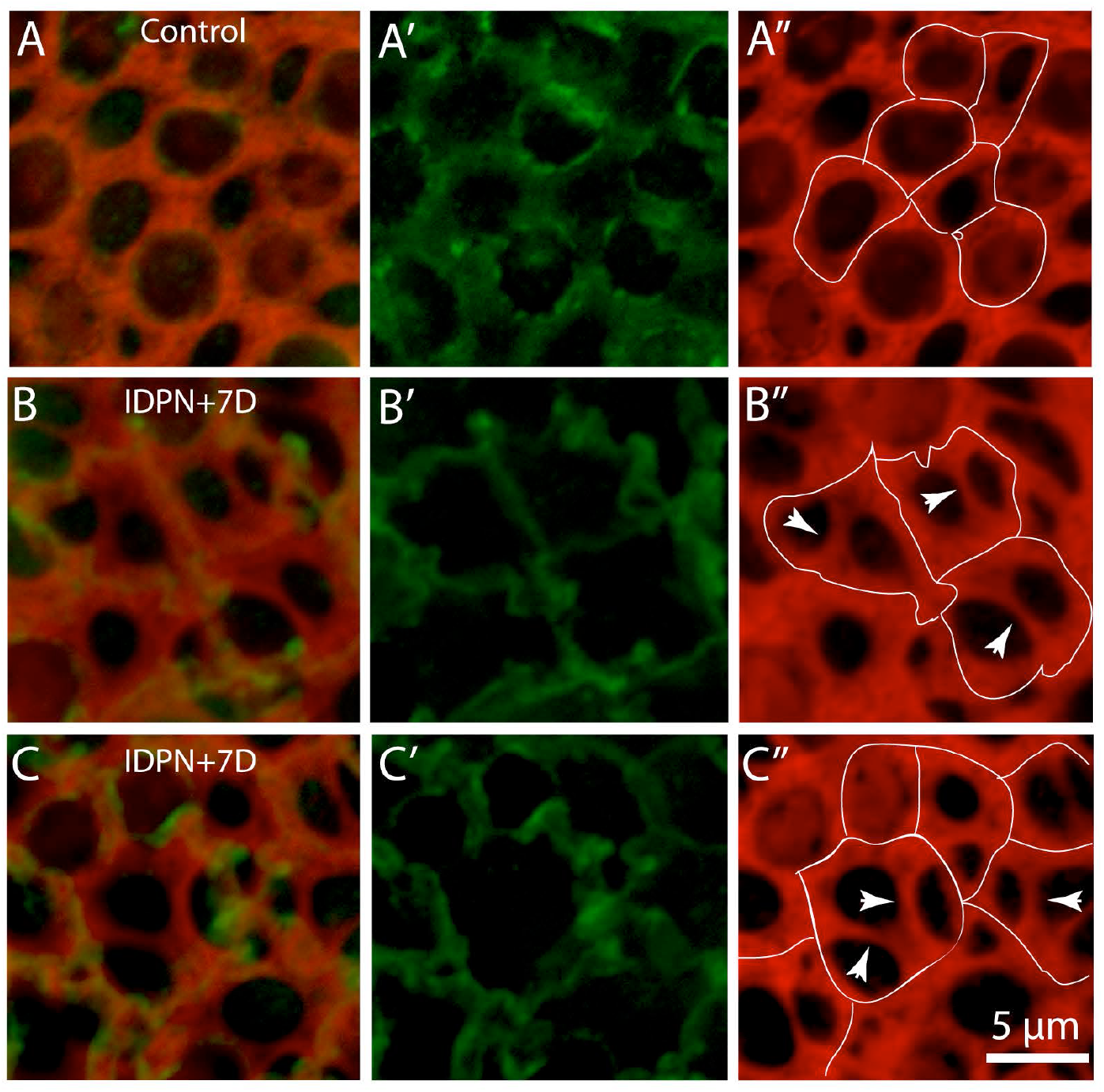
Identification of cell-cell junctions in the mouse utricle after ototoxic lesion. (A): In undamaged utricles, immunolabeling for β-catenin (A, A’, green) overlapped with junctional actin bands (phalloidin, red). Traces of β-catenin immunoreactivity (dotted lines) are shown in A”. (B, C, D) Damaged utricles possess both thick and thin actin bands, making it difficult to identify the sites of cell junctions. lmmuno reactivity for β-catenin (B’, C’, D’,green) labels adherins junctions. Many actin bands are no longer associ ated with β-catenin (B”,C”D”, arrowheads), suggesting that those bands no longer designate cell-cell junctions.

The number of supporting cells in whole-mounts of the uninjured utricle can be determined by quantifying surfaces bounded by actin cables that do not possess features of hair cells (e.g., stereocilia bundles or myosin VIIa immunoreactivity). The structure of actin bands in chick utricles was not impacted by ototoxic lesion, suggesting that they remained associated with cell-cell junctions. However, as noted above, the lumenal surfaces of injured mouse utricles possessed numerous additional actin bands and the borders of individual cells were difficult to distinguish. It also appeared that the actin bands no longer defined the borders of individual cells. To clarify this issue, we quantified the density of lumenal regions that were bounded by filamentous actin structures, but did not also possess cuticular plates and/or stereocilia bundles (e.g., hair cells). The density of such regions in uninjured utricles was 23.9+3.0 regions/1,000 µm^2^ (n=28 samples from five specimens), which likely reflects the density of supporting cells (Fig. 5A). However, the density of actin-bounded regions in utricles from IDPN-lesioned mice was 66.1+12.1 regions/1,000 µm^2^ (Fig. 5B, C, n=20 samples from five specimens). Since supporting cells of these utricles underwent little (if any) cell division after injury [19], the increase in actin-enclosed surfaces cannot be attributed to cellular addition and does not yield an accurate estimate of supporting cell number. Instead, as noted above, many of the (original) thicker actin cables no longer colocalized with cellular junctions, but were confined to the surfaces of individual supporting cells. The continued presence of these cables, when combined with the thinner actin cables that were formed to mediate epithelial repair, caused a transient increase in actin-bounded regions at the lumen of the epithelium (Fig. 5C). In contrast, the density of actin-bound regions on the surfaces of chick utricles was not affected by ototoxic injury (Fig. 5D-F), suggesting that each actin-enclosed region still reflected an individual cell.

**Figure 5.**
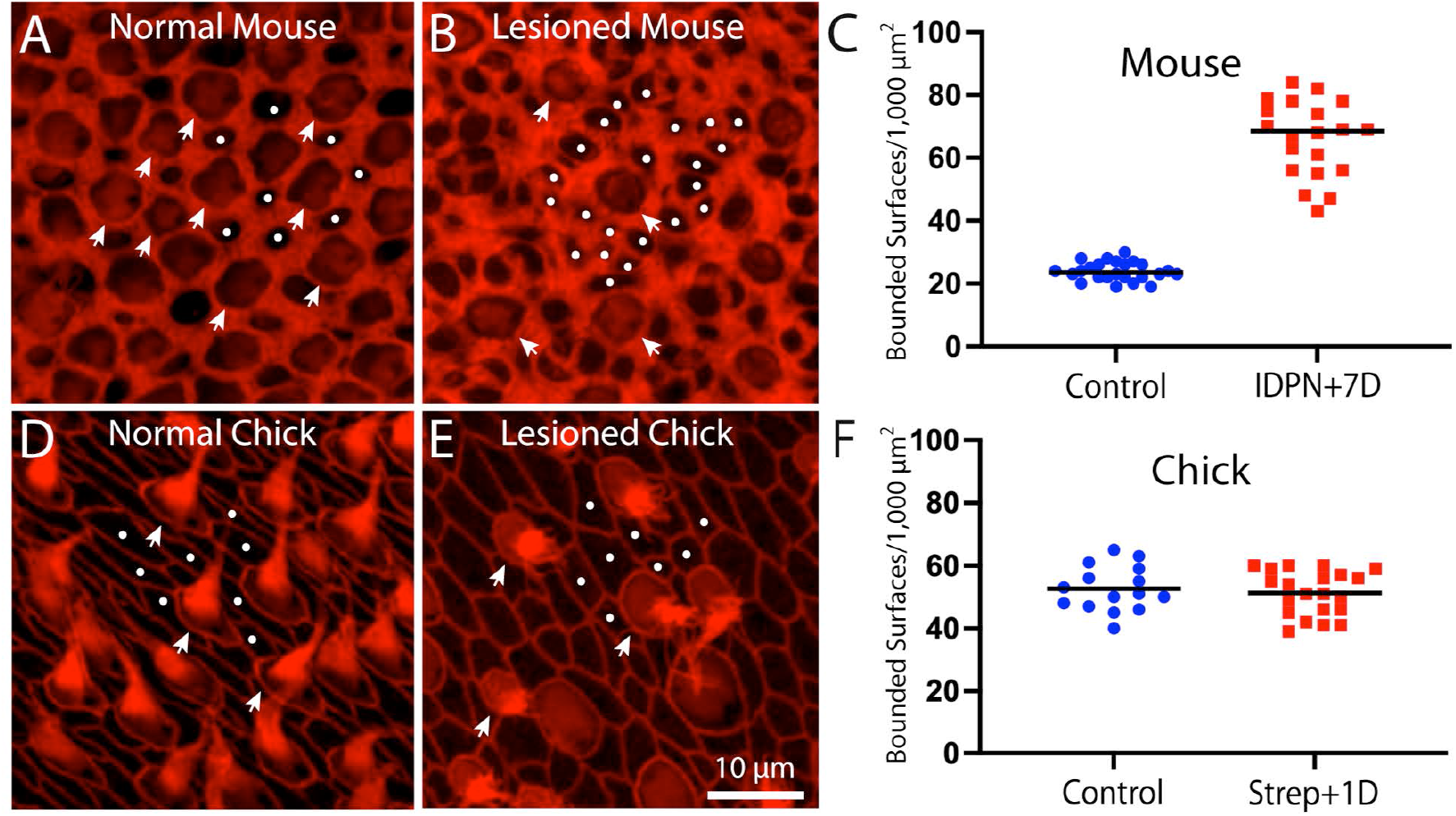
Quantification of F-actin-bound surfaces on the utricles of chicks and mice. (A): Surface view of an uninjured mouse utricle reveals regions that are enclosed by actin bands (phalloidin, red) and can be readily identified as either individual hair cells (arrowheads) or supporting cells (dots). (B): Similar view of a mouse utricle fixed at seven days after IDPN injection reveals many additional actin bound regions (dots), when com pared with the undamaged utricle shown in (A). Also note the presence of fewer hair cells (arrowheads). (C): Quantification of actin-bounded regions without stereocilia or cuticular plates (e.g., dots in A and B) increased by ∼3x after IDPN ototoxicity. (D): Surface view of actin bands in an undamaged chick utricle also permits identi fication of hair cell and supporting cell surfaces. (E): Ototoxic injury to the chick utricle still permits identification of actin-bounded regions that are not hair cells. (F): Quantification of non-hair cell actin-bounded surfaces did not change after ototoxic injury, suggesting that the phalloidin-labeled junctional cables are still associated with cell-cell junctions.

Repair of epithelial lesions will also cause expansion of the surfaces of supporting cells, a process that may trigger a regenerative response. Prior studies have shown that developmental growth and regeneration of the utricle is regulated by the Hippo/YAP signaling pathway, which can be triggered by cellular spreading [11,12,13]. In order to determine the extent of cellular expansion after hair cell loss, we next quantified the surface areas of supporting cells in normal and ototoxically-injured utricles of chicks and mice. Striolar supporting cells in uninjured chick utricles had a mean surface area of 10.8+4.3 µm^2^ (n=140 samples from 5 specimens), while those that had lost ∼40% of their hair cells after streptomycin treatment had a mean surface area of 16.8+8.4 µm^2^ (n=150 samples from 6 specimens; Fig. 6A-C). In contrast, supporting cells in the striolar region of uninjured mouse utricles had a mean surface area of 20.6+7.0 µm^2^, while those that had lost ∼60% of their hair cells after IDPN treatment had a mean surface area of 26.5+8.7 µm^2^ (Fig. 6D-F; n=121/110 samples from six specimens/group). These data reveal some important differences between supporting cells in the utricles of chicks and mice. First, supporting cells in uninjured mouse utricles have surface areas that are about twice as large as those in uninjured chick utricles (Fig. 6C, F). In addition, the loss of ∼40% of hair cells from the chick utricle caused supporting cells to expand by ∼60%, while a greater degree of hair cell loss in mouse utricles (∼60%) led to expansion of supporting cell surfaces by only ∼25%. In order rule out any possible influence of species-specific differences in the apical surfaces of hair cells, we quantified surface areas from hair cells in the striolar region of utricles from chicks and mice. Chick hair cells had a mean surface area of 16.7+7.1 µm^2^ (n=158 samples from five specimens), while mouse hair cells had a mean surface area of 17.1+4.6 µm^2^ (n=150 samples from five specimens). These data suggest that the differences in injury-evoked supporting cell expansion in chick vs. mouse cannot be attributed to differences in the surface areas of (lost) hair cells. In addition, hair cell density in uninjured utricles of mice and chicks is very similar (compare data in Fig. 1F and Fig. 3C). Our observations indicate that the surface areas of supporting cells in normal chick utricles are smaller than those in normal mouse utricles, and that chick supporting cells undergo a greater relative degree of cellular expansion in response to hair cell injury than do supporting cells of the mouse utricle. As discussed below, these observations have implications for differential activation of Hippo/YAP signaling in these two structures.

**Figure 6.**
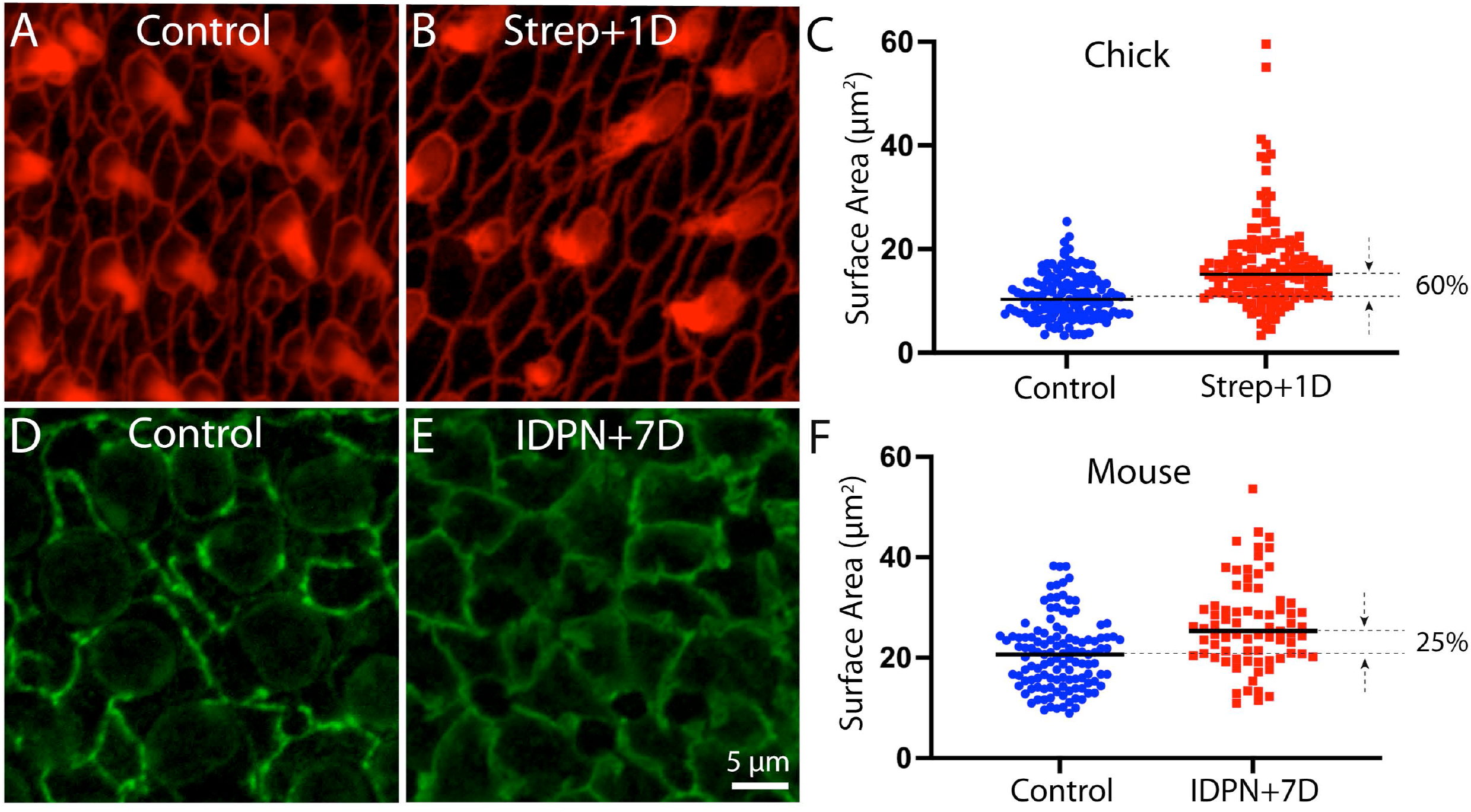
Quantification of the surface areas of supporting cells in normal and ototoxically-lesioned utricles. (A, B): Surface images of the chick utricle after labeling with phalloidin (red). (C): Although the number of supporting cell surfaces does not change after lesion (see Fig. 5), indi vidual supporting cells expand their surfaces by ∼60%, in order to restore the integrity of the epithelium. (D, E): Cell-cell junctions in the mouse utricle were labeled with -catenin (green), which permitted accurate quantification of cell surface area. (F): Resulting data indicate that loss of hair cells caused mouse supporting cells to expand by only ∼25%, which is sufficient to repair the epithelium. The different relative expansion of chick vs. mouse supporting cells can be attributed to the differing numbers and sizes of these cells in both species (see text).

## Discussion

### Link between actin junctions, cell shape and regenerative proliferation

The vestibular organs of nonmammalian vertebrates are capable of regenerating hair cells after ototoxic injury [23]. Most replacement hair cells are produced by the proliferation of supporting cells, which divide to yield new hair cells and supporting cells. In contrast, hair cell regeneration in the vestibular organs of mammals is very limited and occurs by a process of transdifferentiation, whereby supporting cells change phenotype into replacement hair cells without first undergoing cell division [6]. Importantly, these newly-generated hair cells do not acquire mature physiological properties and do not restore proper vestibular function [3,4].

The signals that regulate cell proliferation in the vertebrate inner ear are not fully understood, but a series of studies has suggested that the thickness of circumferential actin belts at cell-cell junctions may influence the potential for cell cycle entry [15]. Cellular junctions in the chick utricle possess thin actin cables, that permit significant cellular spreading and proliferation. In contrast, the junctional actin belts in the mouse utricle are considerably thicker than their chick counterparts, and this appears to diminish the ability of isolated sensory epithelia to spread on defined substrates [7]. Also, during the development of the mouse utricle, the gradual thickening of cellular junctions is correlated with reduced proliferative ability [9]. Finally, the thick circumferential actin bands of the mouse utricle result in high epithelial stiffness, when compared to the chick utricle [24]. The present results demonstrate that ototoxic injury to the mouse utricle results in the formation numerous thin actin bands, many of which are of comparable thickness to those in the chick utricle (Fig. 3D). It is likely that many of these thin bands are created shortly after hair cell loss, in order to facilitate epithelial repair. It would be of interest to quantify the mechanical properties of acutely-injured vestibular epithelia of mice and resolve how the combination of thin and thick actin bands impacts overall epithelial stiffness.

The YAP signaling pathway has been shown to regulate cell proliferation in the utricles of chicks and mice [11,12,13]. Although the mechanisms responsible for YAP activation in the injured utricle are not completely understood, Hippo/YAP signaling can be activated via mechanical stretching [25]. Activation of YAP signaling (as determined by nuclear translocation of the YAP1 protein) occurs in the injured chick utricle, but not in the injured mouse utricle [12,13], so it is notable that the relative degree of injury-induced cellular expansion differs in the utricles of chicks vs. mice. Partial hair cell loss in the chick utricle (∼40%) caused nearby supporting cells to expand their surfaces by about 60%, while a greater degree of hair cell loss in the mouse utricle (∼60%) led to a ∼25% expansion of supporting cells. This process is illustrated schematically in Fig. 7. In many cell types, mechanical stress (such as that caused by cellular expansion) inhibits Lats1/2, thereby preventing the degradation of YAP1 and allowing it to enter the nucleus and activate Hippo pathway genes [27]. The observation that treatment with a small molecule inhibitor of Lats kinases (an upstream regulator of YAP) can induce nuclear translocation of YAP1 in supporting cells of the mouse utricle [26] suggests that signals upstream of Lats1/2 limit the proliferative ability of supporting cells. As such, it is possible that the extent of expansion that occurs in mouse supporting cells after hair cell injury may be insufficient to inhibit Lats kinases, thereby preventing YAP1 translocation and regenerative proliferation.

**Figure 7.**
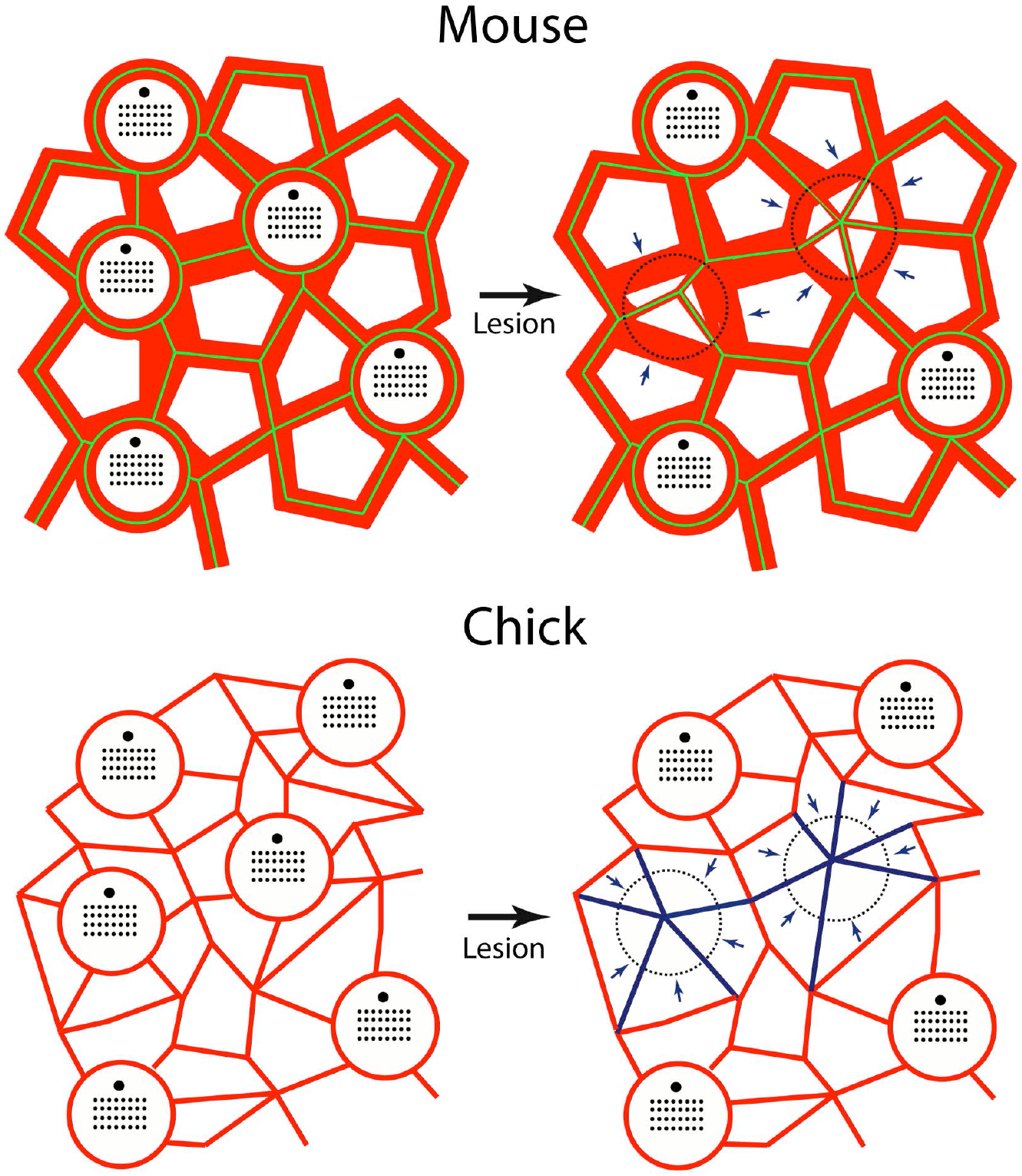
Mechanisms of epithelial repair in the utricles of mice and chicks. Junctional actin bands (red) in the mouse utricle are much thicker than those in the chick. Adherins junctions in the uninjured mouse utricle (green, based on β -catenin immunoreactivity-see Fig. 6) colocalize with these junctional bands. After hair cell loss, supporting cells extend thin actin bands to repair the epithelium and these also form new adherins junctions. Note that many of the remaining thick bands no longer possess adherins junctions (arrows), but are likely to remain within individual supporting cells. Cell-cell junctions in the chick utricle contain thinner actin bands, which rearrange to close the sensory epithelium after hair cell injury (arrows, blue). Also note that the number of supporting cell surfaces in the undamaged chick utricle is about twice that of the mouse utricle. For this reason, hair cell loss causes the surface area of chick supporting cells to expand by a greater extent than in the mouse utricle.

What factors govern the expansion of supporting cells in response to hair cell loss? Our results indicate that the lumenal surface areas of supporting cells in the mouse utricle are about twice as large as in those in the chick utricle (see Fig. 6). Utricles of mature chicks contain ∼29,000 hair cells and the ratio of hair cells to supporting cells is estimated to be about 1:2.5 [28]. In contrast, the mature mouse utricle contains ∼3,600 hair cells [37] and a hair cell to supporting cell ratio of 1:1.7 [29]. The larger surface areas of mouse supporting cells can probably be attributed to the fact that there are only about half as many supporting cells per hair cell in the mouse utricle than in the chick utricle. Cell-cell junctions at the lumen of the sensory epithelium in utricles of both mice and chicks are quickly repaired after injury, in order to restore the barrier function of the epithelium. Given the smaller surface areas of chick supporting cells, those cells will have to undergo a greater relative degree of expansion during epithelial repair, and this may account (in part) for differences in YAP activation and regenerative proliferation that are observed in the utricles of chicks vs. mice.

### Different methods of ototoxic injury

One limitation of the present study is that different ototoxins were used to damage the ears of mice vs. chicks. These experiments were designed to compare injury response in the ears of mice and chicks *in vivo* following a moderate degree of hair cell loss, but there are currently no regimens that are known to reliably induce ototoxic injury in the ears of both mammals and birds. Vestibular hair cells in mature mice can be lesioned using Pou4f3-huDTR transgenic models [6,16], but prior studies indicate that the mouse vestibular organs are relatively insensitive to aminoglycoside antibiotics (except when applied directly into the inner ear [30]). The vestibular organs of chicks can be readily lesioned by systemic treatment with various aminoglycosides [1,24], but IDPN ototoxicity has not been characterized in nonmammalian species. The ototoxicity IDPN has been well-studied in rodents [22] and a significant hair cell lesion can be induced from a single systemic injection (e.g., Fig. 1). The extent of hair cell regeneration that is observed in mice after IDPN treatment is comparable to that observed after diphtheria toxin administration in the Pou4f3-huDTR model [6,23]. Since regenerated hair cells are generated by supporting cells, the fact that regeneration occurs after IDPN ototoxicity suggests that IDPN does not have any deleterious effects on mouse supporting cells. Similarly, we found that the utricles of both chicks quickly resealed their epithelial barriers (via supporting cell extension), despite be treated with differing ototoxins. The similar extent of the ototoxic lesions (∼60% in the mouse utricle, ∼40% in the striola of the chick utricle), combined with rapid epithelial repair in both species, suggests that the injury responses were similar in both cases. Still, we acknowledge that a more optimal approach would involve use of the same ototoxin (and at similar doses) in both chick and mouse.

## Acknowledgments

I thank Jaci Lett for mouse breeding and genotyping. Research was supported by grant R01DC006283 from the NIDCD (NIH)

## Statements and Declarations

The author has no financial interests or conflicts that are relevant to this study. Data will be made available upon reasonable request.

